# Manifold learning analysis suggests novel strategies for aligning single-cell multi-modalities and revealing functional genomics for neuronal electrophysiology

**DOI:** 10.1101/2020.12.03.410555

**Authors:** Jiawei Huang, Jie Sheng, Daifeng Wang

## Abstract

Recent single-cell multi-modal data reveal multi-scale characteristics of single cells, such as transcriptomics, morphology, and electrophysiology. However, our understanding of functional genomics and gene regulation leading to various cellular characteristics remains elusive. To address this, we applied multiple machine learning methods to align gene expression and electrophysiological data of single neuronal cells in the mouse brain. We found that nonlinear manifold learning outperforms other methods. After manifold alignment, the cell clusters highly correspond to transcriptomic and morphological cell-types, suggesting a strong nonlinear relationship between gene expression and electrophysiology at the cell-type level. The aligned cells form developmental trajectories and show continuous changes of electrophysiological features, implying the underlying developmental process. We also found that the manifold-aligned cell clusters’ differentially expressed genes can predict many electrophysiological features. Functional enrichment and gene regulatory network analyses for those cell clusters revealed potential genome functions and molecular mechanisms from gene expression to neuronal electrophysiology.

## Introduction

Recent single-cell technologies have generated a great deal of excitement and interest in studying functional genomics at cellular resolution^1^. For example, recent Patch-seq techniques enable measuring multiple characteristics of individual neuronal cells, including transcriptomics, morphology, and electrophysiology in the complex brains, also known as single-cell multi-modal data^2^. Further computational analyses have clustered cells into many cell types for each modality. The same type’s cells share similar characteristics: t-type by transcriptomics and etype by electrophysiology. Those cell types build a foundation for uncovering cellular functions, structures, and behaviors at different scales. For instance, previous correlation-based analyses found individual genes whose expression levels linearly correlate with electrophysiological features in excitatory and inhibitory neurons^3,4^. Besides, recent studies have also identified several cell types from different modalities that share many cells (e.g., me-type), suggesting the linkages across modalities in these cells^2,5^. However, understanding the molecular mechanisms underlying multi-modalities that typically involve multiple genes is still challenging.

Transcriptomic activities such as gene expression for cellular characteristics and behaviors are fundamentally governed by gene regulatory networks (GRNs)^6^. In particular, the regulatory factors (e.g., transcription factors) in GRNs work together and control the expression of their target genes. Also, GRNs can be inferred from transcriptomic data and be employed as robust systems to infer genomic functions^7^. Many computational methods have been developed to predict the transcriptomic cell-type GRNs using single-cell genomic data such as scRNA-seq^6^. Primarily, relatively little is known about how genes function and work together in GRNs to drive cross-modal cellular characteristics (e.g., from t-type to e-type).

Further, integrating and analyzing heterogeneous, large-scale single-cell datasets remains challenging. Machine learning has emerged as a powerful tool for single-cell data analysis, such as t-SNE^8^, UMAP^9^, and scPred^10^, to identify transcriptomic cell types. An autoencoder model has recently been used to classify cell types using multi-modal data^11^. However, these studies were limited to building an accurate model as a “black box” and lacked any biological interpretability from the box, especially for understanding functional genomics in various cellular phenotypes. To address this challenge, we applied manifold learning, an emerging machine learning field, to align single-cell gene expression and electrophysiological data in the multiple regions of the mouse brain. We found that the nonlinear manifold alignment outperforms other methods for aligning cells from multi-modalities. Also, it identified biologically meaningful cross-modal cell clusters on the latent spaces after the alignment. This suggests a strong nonlinear relationship (manifold structure) linking genes and electrophysiological features at the cell-type level. The aligned cells by manifold alignment show strong developmental trajectories, suggesting the underlying neuronal development and continuous changes of several electrophysiological features. We further found that many electrophysiological features can be predicted by differentially expressed genes of cross-modal cell clusters. Our enrichment analyses for the cell clusters, including GO terms, KEGG pathways, and gene regulatory networks, further revealed the underlying functions and mechanisms from genes to cellular electrophysiology in the mouse brain.

## Results

We have applied multiple machine learning methods to align the single cells in the mouse brain using their gene expression and electrophysiological data (Methods, Fig. 1A). In particular, we focused on two major brain regions, mouse visual cortex and motor cortex, and used the latest Patch-seq data from Allen Brain Atlas in the BRAIN Initiative^5,12,13^ (Methods). The machine learning methods for alignment include linear manifold alignment (LM), nonlinear manifold alignment (NMA), manifold warping (MW), Canonical Correlation Analysis (CCA), Principal Component Analysis (PCA, no alignment) and t-Distributed Stochastic Neighbor Embedding (t-SNE, no alignment). We found that NMA better aligns cells in both regions than other methods, and also uncovers the developmental trajectories of the aligned cells.

**Figure 1.**
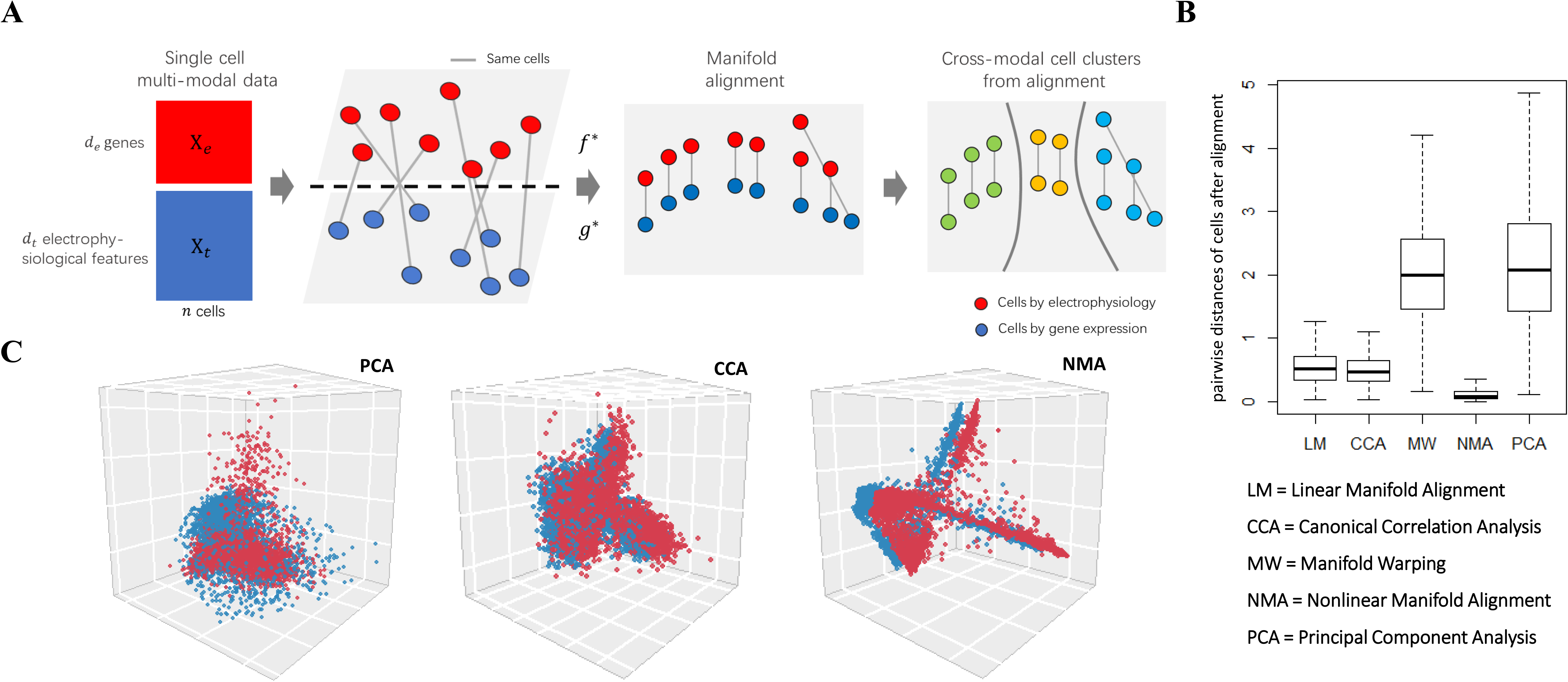
Manifold learning aligns single-cell multi-modal data and reveals nonlinear relationships between cellular transcriptomics and electrophysiology. (**A**) Manifold learning analysis inputs single-cell multi-modal data: *X_e_*, the electrophysiological data (red, *d_e_* electrophysiological features by *n* cells) and *X_t_*, the gene expression data (blue, *d_t_* genes by *n* cells). It then aims to find the optimal functions *f*^*^(.) and *g*^*^(.) to project *X_e_* and *X_t_* onto the same latent space with dimension *d*. Thus, it reduces the dimensions of multi-modal data of n single cells to 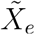 (*d* reduced electrophysiological features by *n* cells) and 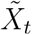 (*d* reduced gene expression features by *n* cells). If manifold learning is used, then the latent space aims to preserve the manifold structures among cells from each modality, i.e., manifold alignment. Finally, it clusters the cells on the latent space to identify cross-modal cell clusters. (**B**) Boxplots show the pairwise cell distance (Euclidean Distance) after alignment on the latent space for 3654 neuronal cells (aspiny) in the mouse visual cortex (Methods). The cell coordinates on the latent space are standardized per cell (i.e., each row of 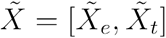) to compare methods. Each box represents one alignment method. The box indicates the lower and upper quantiles of the data, with a horizontal line at the median. The vertical line extended from the boxplot shows a 1.5 interquartile range beyond the 75th percentile or 25th percentile. The machine learning methods for alignment include linear manifold alignment (LM), nonlinear manifold alignment (NMA), manifold warping (MW), Canonical Correlation Analysis (CCA), and Principal Component Analysis (PCA, no alignment). (**C**) The cells on the latent space (3D) after alignment by PCA (no alignment), CCA, and NMA. (**D**). The red and blue dots represent the cells from gene expression and electrophysiological data, respectively. The blue dots are drifted −0.05 on the y-axis to show the alignment.

### Manifold learning aligns single-cell multi-modal data and reveals nonlinear relationships between cellular transcriptomics and electrophysiology

For the visual cortex, after data processing and feature selection (Methods), we aligned 3654 neuronal cells (aspiny) in the mouse visual cortex using their gene expression and electrophysiological data of single cells by Patch-seq. After alignment, we projected the cells onto a low dimensional latent space and then clustered them into multiple cell clusters. The cells clustered together imply that they share both similar gene expression and electrophysiological features. We found that nonlinear manifold alignment outperforms other methods (Fig. 1B) based on the Euclidean distances of the same cells on the latent space. This result suggests potential nonlinear relationships between the transcriptomics and electrophysiology in those neuronal cells, better identified by manifolds. Finally, we visualized the cell alignments of NMA, CCA, and PCA on the 3D latent space in Fig. 1C, showing that nonlinear machine learning has the best alignment (average distances of aligned same cells: PCA = 2.117, CCA = 0.510, NMA = 0.132). In addition, we applied our analysis to another multi-modal data of 102 neuronal cells in the mouse visual cortex and also found that the nonlinear manifold alignment outperforms other methods (Fig. S1). Also, for the motor cortex, after aligning its 1208 neuronal cells using the gene expression and electrophysiological features (Methods), we found a similar result that the NMA outperforms other methods in terms of alignment (Fig. S2, average distances of aligned same cells: PCA = 2.117, CCA = 0.510, NMA = 0.132).

### Manifold-aligned cells recover known cell types and also uncover developmental trajectories across transcriptomic types and continuous changes of electrophysiological features

After aligning single cells using multi-modal data, we found that the aligned cells on the latent space by manifold learning recovered the known cell types of a single modality. For instance, those neuronal cells were previously classified into six major transcriptomic types (t-types) based on the expression of marker genes. We also found that the t-types are better formed and recovered by the latent space of NMA than other methods (e.g., CCA and PCA) (Fig. 2A, Fig. S2) in both regions. In particular, using the t-types of the cells, we calculated the cells’ silhouette values on the latent space after alignment to quantify how well the coordinates of the aligned cells correspond to the t-types (Methods). We found that the silhouette values of NMA are significantly larger than other methods (Fig. 2B), suggesting that NMA better recovers the t-types.

**Figure 2.**
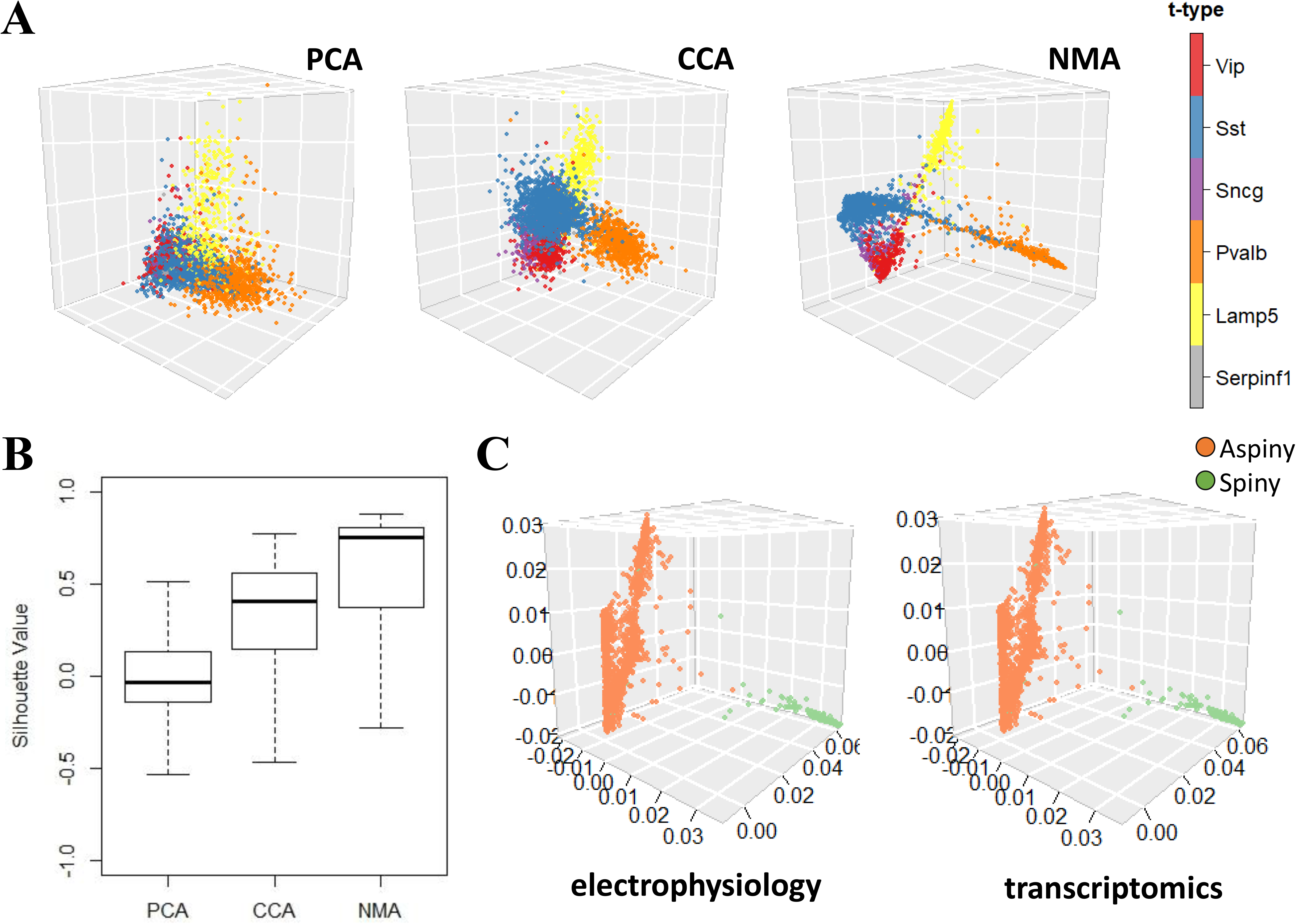
Manifold alignment of single cell multi-modalities recovers known cell types. (**A**) Scatterplots show 3645 neuronal cells in the mouse visual cortex from electrophysiological data on the latent spaces (3D) after alignment by PCA (no alignment), CCA, and NMA. The cells are colored by prior known transcriptomic types (t-types). Red: Vip type; Blue: Sst type; Purple: Sncg type; Orange: Pvalb type; Yellow: Lamp5 type; Gray: Serpinf1 type. The cells from gene expression data on the latent spaces were shown in supplement Fig 3. (**B**) The box plots show the silhouette values of cells for quantifying how well the coordinates of the cells on the latent spaces correspond to the t-types by PCA, CCA and NMA (Methods). (**C**) Scatterplots show neuronal cells in the mouse visual cortex on the latent spaces (3D) after alignment by NMA. Dots are colored according to the reconstructed morphological types (orange: aspiny, lightgreen: spiny).

Also, NMA revealed a pseudo-timing order across these t-types in the visual cortex, implying potential neuronal development aligning with cellular electrophysiology. This developmental trajectory (from Lamp5 to Vip to Serpinf1/Sncg to Sst to Pvalb) was also supported by previous studies^14^. However, other methods, including CCA, PCA, t-SNE/UMAP^5,13^ as well as recent coupled autoencoder method^11^ do not show either multiple t-types or trajectories across t-types (Fig. 2A, Fig. S2). Besides t-types, the aligned cells by NMA also revealed morphological types, as shown by aspiny vs. spiny cells in Fig. 2C. Thus, these results demonstrate that manifold learning has uncovered known multi-modal cell types from cell alignment. In addition, after using NMA to align cells in the motor cortex, we observed this similar trajectory (Lamp5 to Vip to Sncg to Sst to Pvalb) (Fig. 3A).

**Figure 3.**
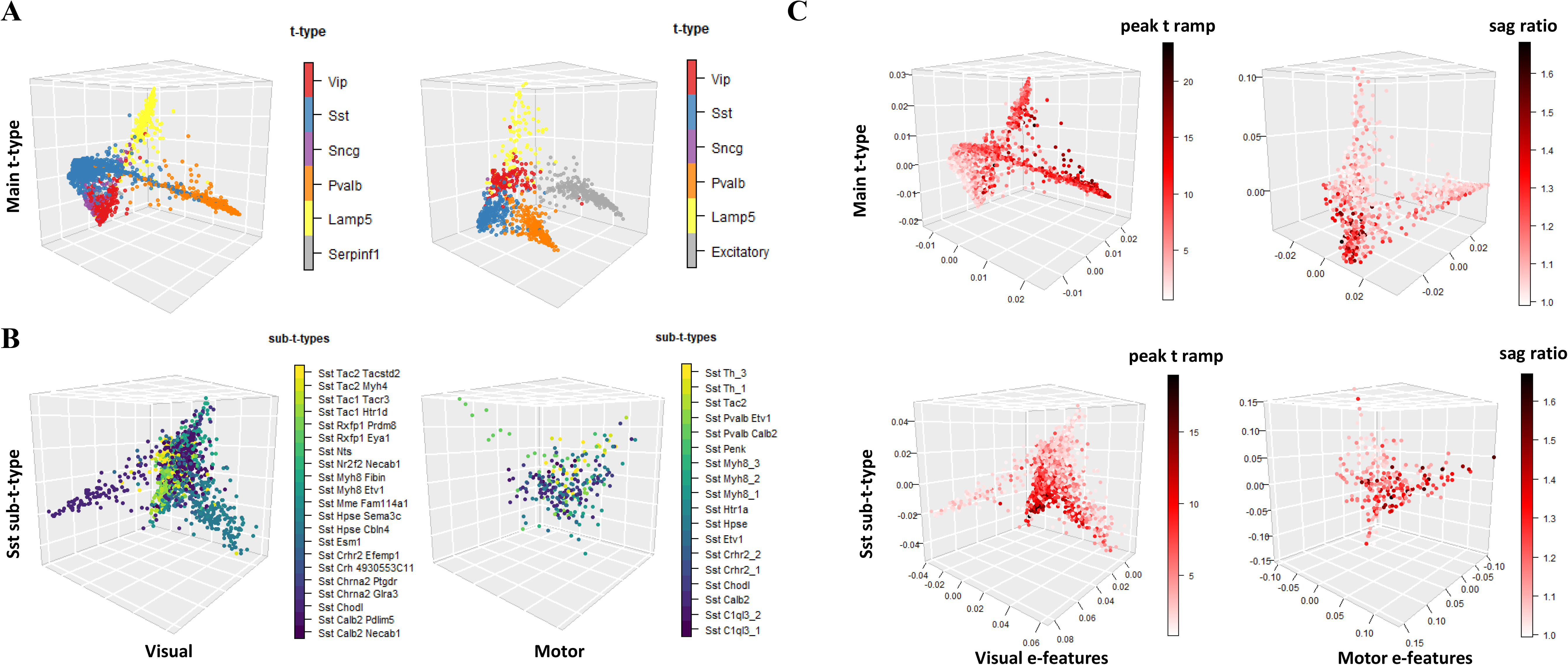
Developmental trajectories across t-types on the latent space after nonlinear manifold alignment along with continuous electrophysiology changes. (**A**) Scatterplots show developmental trajectories across t-types on the latent space (3D used here) after nonlinear manifold alignment of the cells in the visual cortex (left) and motor cortex (right). Cells in shared t-types between two regions are highlighted with the same color for comparison. Red: Vip type; Blue: Sst type; Purple: Sncg type; Orange: Pvalb type; Yellow: Lamp5 type; Gray: other t-types not shared (e.g., Excitatory neurons in the visual cortex). (**B**) Scatterplots show developmental trajectories for Sst sub t-types on the latent space (3D used here) after nonlinear manifold alignment of the cells in the visual cortex (left) and motor cortex (right). (**C**) Scatterplots show continuous changes of select electrophysiological features in t-types and Sst sub-t-types in the visual cortex (left) and motor cortex (right). The “peak t ramp” is the time taken from membrane potential to AP peak for ramp stimulus.

Using NMA, we also observed similar developmental trajectories in sub-t-types. For instance, the Sst sub-t-type is known to have multi-modal diversity in Layer 5^5^. We also found that the aligned cells by NMA show a developmental trajectory (tac2 to myh8 to hpse to crhr2 to chodl to calb2) in both visual and motor cortices (Fig. 3B). This result suggests the great potential of NMA for revealing the underlying expression dynamics in the sub-t-types. Also, we found that certain electrophysiological features of cells on these trajectories show continuous changes, which are also region-specific. For instance, peak_t_ramp (time taken from membrane potential to AP peak for ramp stimulus) gradually changes from low to high along with the trajectory across both t-types and sst sub-t-types in the visual cortex only, whereas the sag ratio changes from low to high in the motor cortex (Fig. 3C). Also, membrane time and AP amplitude achieve high values in the middle of the trajectory across t-types in the motor cortex only (Fig. S3). These electrophysiological features’ continuous changes imply the region-specific activities, although both regions share similar transcriptomic trajectories.

### Cross-modal cell clusters by manifold alignment reveal genomic functions and gene regulatory networks for neuronal electrophysiology

Furthermore, we want to systematically understand underlying functional genomics and molecular mechanisms for cellular electrophysiology using aligned cells. To this end, we clustered aligned cells on the latent space of NMA without using any prior cell-type information. In particular, we used the gaussian mixture model (GMM) to obtain five cell clusters with optimal BIC criterion (Methods, Fig. S4) in the mouse visual cortex. Those cell clusters are cross-model clusters since they are formed after aligning their gene expression and electrophysiological data. As expected, they are highly in accordance with t-types (Fig. S5). For example, Cluster 4 has ~83.3% Lamp5-type cells (373/448 cells), Cluster 2 has ~77.6% Pvalb-type cells (558/719 cells), Cluster 3 has ~86.6% Sst-type cells (1339/1546 cells) and Cluster 1 has ~79.1% Vip cells (541/684 cells). Besides, Clusters 1 and 5 include ~55.8% Serpinf1 cells (24/43) and ~60.7% Sncg cells (84/214), respectively. Also, we identified differentially expressed genes (DEGs) with adjusted p-value <0.01 as marker genes of cross-modal cell clusters (Fig. 4A, Supplemental File 1). In total, there are 182, 243, 175, 190, and 13 marker genes in Clusters 1, 2, 3, 4, 5, respectively. These cell-cluster marker genes are also enriched with biological functions and pathways (GO terms) among the genes (Supplemental File 2) (Methods). For example, we found that many neuronal pathways and functions are significantly enriched in DEGs of Cluster 1, such as the ion channel, synaptic and postsynaptic membrane, neurotransmitter, neuroactive ligand receptor, and cell adhesion (adjusted p<0.05, Fig. 4B). Further, we linked top enriched functions and pathways of each cross-modal cell cluster to its representative electrophysiological features (Fig. S6), providing potential novel molecular mechanistic insights for neuronal electrophysiology. Since gene expression is fundamentally controlled by gene regulatory networks (GRNs), we predicted the GRNs for cross-modal clusters, providing mechanistic insights for multi-modal characteristics (Methods). In particular, the predicted GRNs link transcription factors (TFs) to the cluster’s genes (Supplemental File 3), suggesting the gene regulatory mechanisms for the electrophysiological features in each cluster. For instance, we found that several key TFs on neuronal and intellectual development regulate the genes in Cluster 1, such as Tcf12 and Rora (Fig. 4C). Also, Atf3, a TF modulating immune response^15^, is regulated by inflammatory TFs, Irf5 and Spi1 in the gene regulatory networks of our clusters. Although there are cells not expressing some of these genes, due to the potential off-target expression of immunological genes in Patch-seq^16^, many cells still show high and correlated expression of Atf3, Irf5, and Spi1 (Fig. S7). This observation thus suggests potential interactions between neurotransmission and inflammation, which were recently reported^17^. Besides, Lhx6, a TF previously found inducing Pvalb and Sst neurons^18^, was also predicted as a key TF for the Cluster 2 and Cluster 3 only that have most Pvalb and Sst type neurons, respectively. For the motor cortex, we also identified five major cell clusters from the NMA’s latent space. Like the visual cortex, the motor cortex’s cell clusters also correspond to the transcriptomic types (Fig. S5). For instance, Cluster 5 has ~95.4% Vip type cells (146/153 cells). Cluster 4 has ~75.3% Sst type cells (202/271). Besides, Clusters 1 and 3 respectively include ~34.9% (101/289 cells) and ~64.7% (187/289) Pvalb type cells. For excitatory neurons, ~55.8% (218/391) of cells are in Cluster 2, and ~43.7% (171/391) of cells are in Cluster 1. The predicted GRNs for these cell clusters in the motor cortex also reveal key neuronal TFs such as Lhx6 again, Atf4 as stress-inducible TF, and Npdc1 for neural proliferation, differentiation, and control (Supplement File 3). Finally, we also predicted GRNs for known t-types in both regions (Supplement File 4), which, however, do not include several key TFs such as Lhx6 for Pvalb and Sst types.

**Figure 4.**
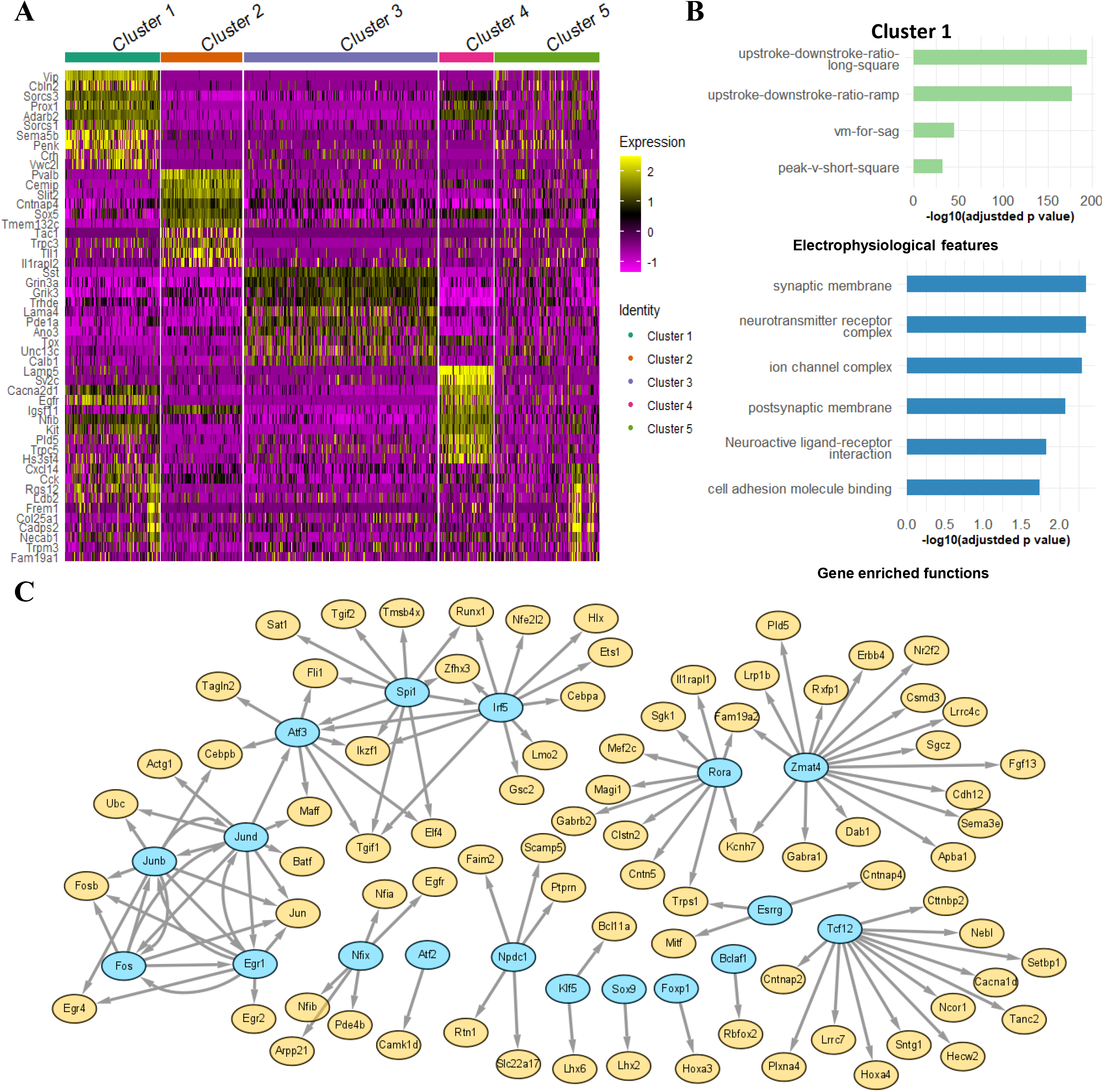
Differentially expressed genes, enrichments, and gene regulatory networks for cross-modal cell clusters. (**A**) The gene expression levels across all 3654 cells for Top 10 differential expressed genes (DEGs) of each cross-modal cell cluster in the mouse visual cortex. The cell clusters were identified by the gaussian mixture model (Methods). The expression levels are normalized (Methods). (**B**) The select enriched biological functions and pathways of DEGs (GO and KEGG terms with adjusted p-value <0.05) and representative electrophysiological features (adjusted p-value <0.05) in Cluster 1 of the mouse visual cortex. (**C**) Gene regulatory networks that link transcription factors (TFs, cyan) to target genes (Orange) in Cluster 1 of the mouse visual cortex.

### Predicting electrophysiological features from gene expression using manifold alignment results

Finally, we want to see if the electrophysiological features could be predicted by gene expression using our manifold alignment. First, we visualized the NMA’s latent spaces of the cells using the bibiplot method (a group of biplots)^5^ (Fig. 5A for the visual cortex and Fig. 5B for the motor cortex). In particular, we selected the first three components of transcriptomic space and electrophysiological space so that each biplot shows such a space using two components. Due to the nonlinear manifold alignments, the transcriptomic spaces and electrophysiological spaces look much more similar than previously used linear dimensionality reduction^5^. This suggests that in general, transcriptomics and electrophysiology have strong nonlinear associations. As shown in each biplot, a group of highly correlated genes and electrophysiological features with the NMA’s latent spaces are highlighted by lines (the line length, i.e., radius, corresponds to the correlation value with max correlation = 1). We found that many genes and electrophysiological features are in similar directions in the biplots, suggesting their strong associations on the NMA’s latent space. For instance, peak_t_ramp and Pavlb are in similar directions on the first and second component of the visual cortex (Fig. 5A), and peak_t_ramp indeed has high values in the Cluster 2 that is enriched with Pvalb cells (Fig. 3C, Fig. S6). Furthermore, we applied a multivariate regression model to fit the components of the NMA’s electrophysiological space (dependent variables) by the components of the NMA’s latent transcriptomic space (independent variables) (Methods). We obtained cross-validated *R*^2^ = 0.987 and *R*^2^ = 0.977 for the visual and motor cortices, respectively. High *R*^2^ values imply that most of the variance of reduced electrophysical data can be explained by transcriptomic data on the NMA’s latent space. Besides, this *R*^2^ value is not affected by the dimensionality of the NMA’s latent space. It slightly decreased as the dimension increases (from 0.987 to 0.954 for the visual cortex; from 0.977 to 0.952 for the motor cortex when we increased the latent space dimensions from 3 components to 20 components).

**Figure 5.**
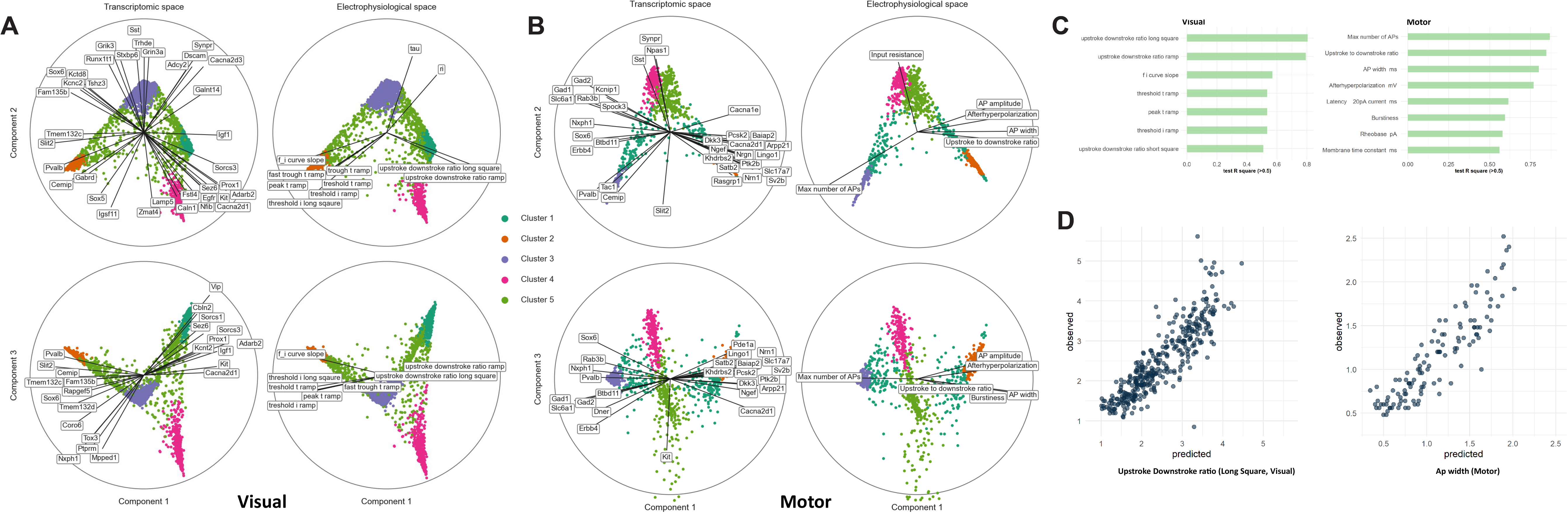
Association and prediction of electrophysiological features from gene expression. (**A**) Bibiplots for the mouse visual cortex using the NMA’s latent spaces (first three components used). Cells are dots (n=3654). Transcriptomic and electrophysiological latent spaces are shown as columns. Each biplot shows the subspace of two components. Cells are colored by their cross-modal clusters. The line length of a gene or electrophysiological feature (i.e., radius) corresponds to its correlation with the latent space with max value = 1. The genes and electrophysiological features with correlations > 0.6 are shown here. The label positions are slightly adjusted to avoid overlapping. (**B**) Similar to (**A**) but for the mouse motor cortex (n=1208). (**C**) The representative electrophysiological features in the cross-modal clusters with testing *R*^2^ > 0.5 (90% training set, 10% testing set, see Methods). (**D**) The predicted values by gene expression (x-axis) vs. the observed values (y-axis) of the upstroke downstroke ratio (*R*^2^ = 0.805) in the visual cortex and the action potential width (*R*^2^ = 0.800) in the motor cortex.

Second, after showing strong associations between genes and electrophysiological features on the NMA’s latent space, we next tried to predict the electrophysiological features from gene expression from our cross-modal clusters (Methods). Specifically, we selected the representative electrophysiological features from each cluster as potentially predictable features. We then fitted a linear regression model to predict each representative electrophysiological feature (dependent variable) by the expression levels of differentially expressed genes (adjusted p-value < 0.05) from the same cluster of the feature across all cells. We also split the cells into 90% training and 10% testing sets and calculated the fitting *R*^2^ values of testing sets (Supplement File 5). For example, for Cluster 1 in the mouse visual cortex, we used its 182 differential expressed genes to predict the upstroke downstroke ratios for long square and ramp and achieved *R*^2^ =0.805 and 0.794, respectively. As shown in Fig. 5C, a number of electrophysiological features can be predicted by differential expressed genes with *R*^2^ > 0.5. In addition, Fig. 5D shows that the predicted values are highly correlated with the observed values across many cells for the upstroke downstroke ratio (*R*^2^ = 0.805) in the visual cortex and the action potential width (*R*^2^ = 0.800) in the motor cortex. Moreover, we compared this result with the testing *R*^2^ of predicting electrophysiological features based on the differentially expressed genes of known t-types. In comparison, we obtained an *R*^2^ = 0.765 for predicting the action potential width in the motor cortex and an *R*^2^ = 0.725 for predicting the upstroke downstroke ratio for long square stimulus in the visual cortex. This suggests the great potential of using our cross-modal clusters from nonlinear manifold alignment along with differentially expressed genes for improving predicting electrophysiological features from gene expression.

## Discussion

This study applied manifold learning to integrate and analyze single cells’ gene expression and electrophysiological data in the mouse brain. We found that the cells are well aligned by the two data types and form multiple cell clusters after manifold alignment. These clusters were enriched with neuronal functions and pathways and uncovered additional cellular characteristics, such as morphology and development. Our results suggest the great potential of manifold learning to analyze increasing single-cell multi-omics data and understand single-cell functional genomics in the near future. Our manifold learning analysis is general-purpose and enables studying single-cell multi-modal data in the human brain and other contexts^19^. Moreover, our GRN analysis can also serve as a basis for understanding gene regulation for additional cellular multi-modal phenotypes.

Besides, this work used several electrophysiological features to represent the characteristics of neuronal electrophysiology that likely miss additional information such as continuous dynamic responses to stimulus. Thus, using advanced machine learning methods such as deep learning for time series classification^20^ to directly integrate time-series electrophysiological data with transcriptomic data will potentially reveal deeper relationships across the modalities and improve cell-type classifications. The predicted gene regulatory networks in this study focused on linking transcription factors to target genes on the transcriptomic side. However, gene regulation is a complex process involving many genomic and epigenomic activities such as chromatin interactions and regulatory elements. Thus, integrating emerging single-cell sequencing data such as scHi-C^21^ and scATAC-seq^22^ as additional modalities will help understand gene regulatory mechanisms in cellular characteristics and behaviors.

## Methods

### Single-cell multimodal datasets

We applied our machine learning analysis for multiple single-cell multimodal datasets in the mouse brain.

#### Visual cortex

Primarily, we used a Patch-seq dataset that included the transcriptomic and electrophysiological data of 4435 neuronal cells (GABAergic cortical neurons) in the mouse visual cortex^13^. In particular, the electrophysiological data measured multiple hyperpolarizing and depolarizing current injection stimuli and responses of short (3 ms) current pulses, long (1 s) current steps, and slow (25 pA/s) current ramps. The transcriptomic data measured genomewide gene expression levels of those neuronal cells. Six transcriptomic cell types (t-types) were identified among the cells: Vip, Sst, Sncg, Serpinf1, Pvalb, and Lamp5. Further, morphological information was provided: 4293 aspiny and 142 spiny cells. Also, we tested our analysis for another Patch-seq dataset in the mouse visual cortex^12^. This dataset includes 102 neuronal cells with electrophysiological data and gene expression data (Fig. S1).

#### Motor cortex

Another Patch-Seq dataset that including the transcriptomic and electrophysiological data of 1227 neuronal cells (GABAergic cortical neurons) in the mouse motor cortex^5^. The electrophysiological data measured multiple hyperpolarizing and depolarizing current injection stimuli and responses of long current steps. The transcriptomic data measured genome-wide gene expression levels of those neuronal cells. Five major transcriptomic cell types (t-types) were identified among the cells: Vip, Sst, Sncg, Pvalb, and Lamp5, based on which 90 neuronal sub-t-types were also labeled.

### Data processing and feature selection of multi-modal data

#### Visual cortex

For electrophysiology, we first obtained 47 electrophysiological features (efeatures) on stimuli and responses, which were identified by Allen Software Development Kit (Allen SDK) and IPFX Python package^23^. Second, we eliminated the features with many missing values such as short_through_t and short_through_v as well as the cells with unobserved features, and finally selected 41 features in all three types of stimuli and responses for 3654 aspiny cells (inhibitory) and 118 spiny cells (excitatory) out of the 4435 neuronal cells. Since the spiny cells usually don’t contain the t-type information, we used the 3654 aspiny cells for manifold learning analysis. Together, we used the 3654 aspiny cells and 118 spiny cells to refer to morphological cell types (m-type). Also, we standardized the feature values across all cells to remove potential scaling effects across features for each feature. The final electrophysiological data matrix is *X_e_* (3654 cells by 41 e-features). We selected 1302 neuronal marker genes^24^ and then took the log transformation of their expression levels. The final gene expression data is *X_t_* (3654 cells by 1302 genes).

#### Motor cortex

For electrophysiology, there are 29 electrophysiological features summarized by^5^. We eliminated the cells with missing observations in these features and standardized them across each feature. Then we selected 1208 cells with features aroused by long square stimuli. For gene expression data, we again selected 1329 neuronal marker genes^24^ and then took the log transformation of their expression levels. The final electrophysiological data matrix is *X_e_* (1208 cells by 29 e-features), and the gene expression data is *X_t_* (1208 cells by 1329 genes).

### Manifold learning for aligning single cells using multi-modal data

We applied manifold learning to align single cells using their multimodal data to discover the linkages of genes and electrophysiological features. In particular, the manifold alignment projects the cells from different modalities onto a lower-dimensional common latent space for preserving local nonlinear similarity of cells in each modality (i.e., manifolds). The distances of the same cells on the latent space can quantify the performance of the alignment. Specifically, given *n* single cells, let 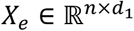 and 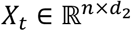 represent their electrophysiological and gene expression data where *d*_1_ is the number of electrophysiological features, and *d*_2_ is the number of genes. Also, 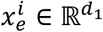 and 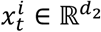 are *i^th^* row of *X_e_* and *X_t_*, representing the electrophysiological and gene expression data of *i^th^* cell. The manifold alignment aims to find optimal projection functions *f*^*^(.) and *g*^*^(.) to map 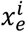, 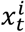 onto a common latent space:

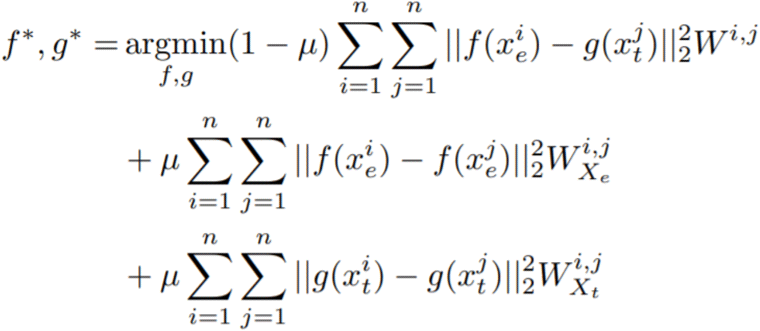

where *f*^*^(.) and *g*\*(.) can be either linear or nonlinear mapping functions, the corresponding matrix 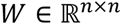 models cross-modal relationships of cells (i.e., identity matrix here), and the similarity matrices *W_X_e__*, 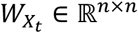 model the relationships of the cells in each modality and can be identified by *k*-nearest neighbor graph (*k*NN). We chose the number of nearest neighbors to be 2, while we also tried other numbers, but the relative performance for different numbers didn’t change much (Fig. S8). The parameter *μ* trades off the contribution between the preserving local similarity for each modality and the correspondence of the cross-modal network. We set *μ* = 0.5. We used our previous ManiNetCluster method^25^ to solve this optimization and found the optimal functions using linear and nonlinear methods, including linear manifold alignment, canonical correlation analysis, linear manifold warping, nonlinear manifold alignment, and nonlinear manifold warping. Finally, after alignment, let 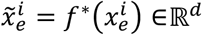 and 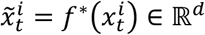 represent the coordinates of the *i_th_* cell on the common latent space (*d*-dimension) and *d* be 3 in our analysis for visualization.

### Identification of cross-modal cell clusters using Gaussian Mixture Model

After alignment, the cells clustered together on the latent space imply that they share similar transcriptomic and electrophysiological features and form cross-modal cell types. To identify such cross-modal cell types, we clustered the cells on the latent space into the cell clusters using gaussian mixture models (GMM) with *K* mixture components. Given a cell, we assigned it to the component *k*_0_ with the maximum posterior probability:

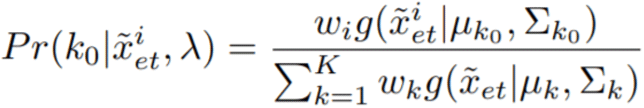

where 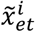 is the *i^th^* row of a combined feature set 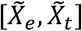, *λ* = {*w_k_, μ_k_*|Σ_*k*_} *k* = 1,…, *K* are parameters: mixture weights, mean vectors, and covariance matrices. Finally, the cells assigned to the same component form a cross-modal cell type. Also, we used the Expectation-maximization algorithm (EM) algorithm with 100 iterations to determine the optimal number of clusters with K=5 (Fig. S4) by Bayesian information criterion (BIC) criterion^26^. K=5 was chosen at which the 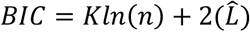 of the model has an approximately constant and insignificant gradient descent through the equation. Silhouette values are used to compare the clustering result^27^, which takes a value from −1 to 1 for each cell and indicates a more pronouncedly clustered cell as the value increases.

### Differentially expressed genes, enrichment analyses, gene regulatory networks, and representative cellular features of cross-modal cell clusters

We used the Seurat to identify differentially expressed genes of each cell cluster and multiple tests, including Wilcox and ROC, to further identify the marker genes of cell clusters (adjusted p-value < 0.01)^28^. We applied this method to the electrophysiological features (absolute values) to find each cluster’s represented e-features. Also, we used the web app, g:Profiler to find the enriched KEGG pathways, GO terms of cell-cluster marker genes, implying underlying biological functions in the cell clusters^29^. Enrichment p-values were adjusted using the Benjamin-Hochberg (B-H) correction. Furthermore, we predicted the gene regulatory networks for cell clusters, linking transcription factors to target marker genes by SCENIC^30^. Those networks provide potentially novel regulatory mechanistic insights for electrophysiology at the cell-type level.

### Prediction of electrophysiological features using gene expression

We generated the bibiplots that consist of a group of biplots using the method in^5^. In particular, we used the first three components of the transcriptomic and electrophysiological latent spaces from nonlinear manifold alignment (NMA) as the latent spaces for generating biplots. For the multivariate linear regression, let 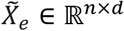 and 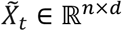 be the first *d* dimensions of the electrophysiological and transcriptomic latent spaces, respectively, where *n* is the number of cells for training. The loss function of the multivariate regression is defined as 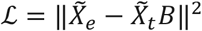, and the solution is given by 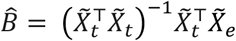. Also, we performed 10-fold cross-validation with 20 repetitions. For each repetition, all cells were randomly partitioned into 10 subsets. A subset was selected as testing set, and remaining subsets were assigned as training sets. The training sets were used to estimate coefficients, and the testing set was used to calculate *R*^2^. The process was repeated 10 times to choose different testing sets. Cross-validated *R*^2^ is calculated through 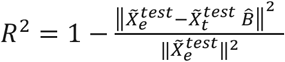, where 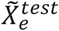 and 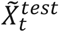 were centered using testing set means. The reported *R*^2^ is averaged across all folds and repetitions. We also tried multiple *d* values to check where the regression overfits, especially for the low dimensionality of the latent space. We varied *d* from 3 to 20 and found that the cross-validated *R*^2^ does not change too much and slightly decreased as the dimension increases (from 0.987 to 0.954 for the visual cortex; from 0.977 to 0.952 for the motor cortex).

Also, we used the multivariate linear regression to predict represented electrophysiological features by the expression levels of differentially expressed genes (DEGs) (adjusted p-value < 0.05) of our cross-modal clusters (and t-types). In particular, we split the cells into 90% training set (*n_traln_* cells) and 10% testing set (*n_test_* cells). Let 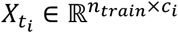 represent the expression levels of *c_i_* differential expressed genes in Cluster *i*, and 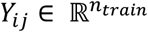 represent the observed values of *j*^th^ represented electrophysiological feature in Cluster *i* for the training cells, the predicted *j*^th^ electrophysiological feature 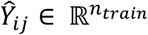 and regression parameters 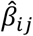 given by 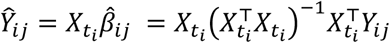, based on the solution to the multivariate linear regression as above. Finally, we can predict the electrophysiological feature for testing set, 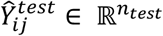 by 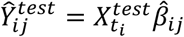, and calculate both training and testing *R*^2^ values for evaluating the prediction.

## Supplementary information

Supplemental materials – Supplemental figures

Supplemental file 1 – Differentially expressed genes in cell clusters

Supplemental file 2 – Enrichments of differentially expressed genes in cell clusters

Supplemental file 3 – Gene regulatory networks for manifold-aligned cell clusters

Supplemental file 4 – Gene regulatory networks for known t-types

Supplemental file 5 – Testing *R*^2^ values for electrophysiological feature prediction from gene expression

## Author contributions

D.W. conceived and designed the study. J.H., J.S. and D.W. analyzed the data and wrote the manuscript. All authors read and approved the final manuscript.

## Acknowledgments

This work was supported by the grants of National Institutes of Health, R01AG067025, R21CA237955 and U01MH116492 to D.W., U54HD090256 to Waisman Center, and the startup funding for D.W. from the Office of the Vice Chancellor for Research and Graduate Education at the University of Wisconsin–Madison.

